# Risk-taking incentives predict aggression heuristics in female gorillas

**DOI:** 10.1101/2025.05.09.653023

**Authors:** Nikolaos Smit, Martha M Robbins

## Abstract

Competition is commonly reflected in aggressive interactions among groupmates, as individuals try to attain or maintain higher social ranks that can offer them better access to critical resources. In this study, we investigate the factors that can shift competitive incentives against higher- or lower-ranking groupmates, that is, more or less powerful individuals. We use a long-term behavioural dataset on five wild groups of the two gorilla species starting in 1998, and we show that most aggression is directed from higher- to lower-ranking adult females close in rank, highlighting rank-reinforcement incentives. Yet, females directed 42% of aggression to higher-ranking females than themselves. Females targeted groupmates of higher rank with increasing number of males in the group, suggesting that males might buffer female-female aggression risk. Contrarily, they targeted females of lower rank with increasing number of females in the group, potentially because this is a low risk option that females prefer when they have access to a larger pool of competitors to choose from. Lactating and pregnant females, especially those in the latest stage of pregnancy, targeted groupmates of higher rank than the groupmates that cycling females targeted, suggesting that energetic needs may motivate females to risk confrontation with more powerful rivals. Our study provides critical insights into the evolution of competitive behaviour, showing that aggression heuristics, the simple rules that animals use to guide their aggressive interactions, are not simply species-specific but also dependent on the conditions that populations and individuals experience.

## 1 Introduction

Animals that live in groups often compete for access to resources such as food and mates [1, 2]. The potential costs of this competition can drive the formation of hierarchies that determine priority of access to resources without superfluous conflicts [3, 4]. Accordingly, individuals may choose strategically who they compete with, in order to minimize costs and maximize gains. Previous research suggests that individuals primarily compete with those closest in the hierarchy, to attain (against close higher-ranking) or maintain (against close lower-ranking individuals) their ranks (‘rank-dependent aggression’; [5–10]). Aggression, probably the most straightforward proxy of competition, commonly increases in frequency when resource availability is lower and it is most often directed towards lower-ranking individuals [8, 11]. Yet, recent research has demonstrated that the rules that animals use to guide their aggressive interactions (‘aggression heuristics’) towards groupmates of different ranks vary even within species [6, 8]. In this study, we test the hypothesis that this variation arises due to the different conditions that (individual) animals experience, and specifically that the social environment and individual energetic needs shape competitive incentives and aggression towards individuals of different ranks.

Regarding the social environment, larger group sizes may entail a larger number of low-ranking individuals which are cumulatively targeted, as competition and aggression are preferentially directed towards the lowest-ranking groupmates (see also ‘bullying’ in [8]). When individuals have access to a larger pool of competitors, they may preferentially target less powerful ones, consistent with risk-sensitivity theory, which posits that individuals tend to minimize risks when they have the option [12, 13]. Conversely, larger group sizes might entail a larger number of allies (e.g., kin for spotted hyaenas, *Crocuta crocuta*, [14]) or protectors (e.g., adult males for female gorillas [11, 15]), which can increase social support and, eventually, minimize any risk-related costs of aggression towards higher-ranking competitors. Hence, group size, the overall number of individuals in the group, might be a poor predictor of aggression because it conflates opposing effects of different kinds of individuals (e.g., see [11] for an example of opposing effects of the number of females and number of males on female gorilla aggression). Finally, if larger group sizes promote within group competition increasing overall aggression rates [2, 11, 16], they might simultaneously promote high-ranking individuals to reinforce their status by targeting lower-ranking groupmates and lower-ranking individuals to direct aggression against higher-ranking groupmates if this can allow them access to precious resources.

Regarding the energetic needs, greater needs for resources may boost the incentives of higher-ranking individuals to reinforce their status by strategically directing their competitive efforts towards lower-ranking groupmates [8, 9, 17]. Contrarily, greater individual needs may also prompt low-ranking individuals, who struggle more to access resources and experience a lower risk-reward ratio (more to gain, less to loose), to show greater aggression rates even against higher-ranking groupmates [18–20]. This aggression may help individuals to improve their ranks but it might be risky if it can incite retaliation from high-ranking, powerful, recipients. Yet, the benefits related to status improvement (long-term benefit) or resource acquisition (short-term benefit) might counterbalance any risk-related costs. Various empirical examples support the latter hypothesis: hunger-driven payoff asymmetries can increase aggression from lower- to higher-ranking individuals (noble crayfish, *Astacus astacus*; [21]), reproductive suppression of low-ranking females can promote conflict escalation against high-ranking ones (paper wasps, *Polistes dominulus*; [22]) and the energetic/nutritional needs of pregnancy (due to support of fetal growth) or lactation (due to milk production) can increase female aggression that reverses hierarchical relationships [23–25].

Studies investigating the influence of social, ecological or physiological factors on aggression patterns across species usually focus on aggression frequency/rate [26, 27]; in this study, we build on this literature to test how relevant factors influence ‘aggression direction’ in terms of power differentials, aiming to unravel another evolutionary aspect of competitive strategies. Gorillas represent an intriguing case for this endeavour, because females of both species form surprisingly stable hierarchical relationships, usually maintained over their whole co-residence in a group, but they often direct aggression towards higher-ranking groupmates [28–30]. Female-female aggression rates towards both higher- and lower-ranking rivals decrease with the number of adult males/protectors in the group but increase with the number of females/competitors in the group [11, 29–31]. Additionally, aggression rates are greater in pregnant females [11] which, like lactating females, spend more time feeding [32], highlighting the greater energetic needs of females in these reproductive states – similar to humans and other apes [33].

We use behavioural observations on one wild western (*Gorilla gorilla gorilla*) and four wild mountain (*Gorilla beringei beringei* ) gorilla groups, starting in 1998 in one of the mountain gorilla groups, to test if the social environment or energetic needs influence female aggression towards more or less powerful females. Specifically, we test whether females direct aggression of lower or higher score (score = recipient-aggressor rank difference) depending on (i) the number of males in their group who may support or protect females, (ii) the number of females in the group representing competitors over resources, and (iii) female reproductive state (cycling, stage of pregnancy, or lactation) which is an (indirect) proxy of energetic needs. We compare our results on the direction of aggression to previous results examining the effects of the same variables on aggression frequency/rates.

## Methods

### Study system and behavioural data

We studied one western gorilla group (ATA/Atanangad; Table 1) in Loango National Park, Gabon, and four mountain gorilla groups in Bwindi Impenetrable National Park, Uganda (Table 1). Observations of western gorillas lasted typically between 07:00 and 16:30h but observations of mountain gorillas were limited to 4 hours per day, typically between 08:00 and 15:00h following the regulations of the Uganda Wildlife Authority.

**Table 1:** Study groups, study period, average number of females per day (±s.d.; total number in parentheses), and number of aggressive interactions (mild/moderate/severe).

| Group | Period | # of females | Aggression |
| --- | --- | --- | --- |
| ATA | 2017-2023 | 3.30±0.85 (6) | 208 / 65 / 27 |
| BIT | 2015-2023 | 4.14±0.35 (6) | 509 / 35 / 23 |
| KYA | 1999-2016 | 6.07±1.02 (9) | 3256 / 445 / 315 |
| MUK | 2016-2023 | 6.59±1.20 (8) | 1389 / 89 / 141 |
| ORU | 2014-2023 | 5.59±0.96 (8) | 355 / 7 / 7 |

Trained observers recorded both focal and ad libitum behavioural observations, including decided avoidance (when an individual walks away from another approaching individual) and displacement (when an individual avoids another and the latter takes the place of the first) behaviours. These two behaviours are ritualized, occurring in absence of aggression, they are considered a more reliable proxy of power relationships over aggression [34], and they are typically used to infer gorilla hierarchical relationships [29, 30]. Similar to the recent studies which have inferred social ranks in gorillas [11, 35], we used all avoidance and displacement interactions throughout the study period and we used the function *elo.seq* from R package *EloRating* [36] to infer daily individual female Elo-scores [37, 38]. We assigned to all avoidance/displacement interactions equal intensity, that is, equal influence to the power relationship of the interacting individuals (k=100). We also assigned to individuals present at the onset of the study initial Elo-scores of 1000 and to individuals entering the hierarchy later (e.g. maturing individuals or immigrants) the score of the lowest ranking individual during the entrance day [11]. This method takes into account the temporal sequence of interactions and updates an individual’s Elo-scores each day the individual interacted with another. The Elo-scores of the winner and loser of an interaction are updated as a function of the winning probabilities prior the interaction: winners with low winning probabilities get greater score increases than winners with high winning probabilities and losers with low winning probabilities get smaller score decreases than losers with high winning probabilities. We present these interactions and hierarchies in detail in [28] (see “traditional Elo rating method”; we do not use the “optimized Elo-rating method” as it yields similar results and it is not widely used). We standardized Elo-scores per group and day such that the highest-score was 1 and the lowest 0.

The observers also recorded aggressive behaviours among adult females (*>*10 years old for western; *>*8 years old for mountain female gorillas [11]), which can be context dependent [34]. For our analysis, we classified these behaviours into three intensity categories (as per [30, 39]): mild (cough/pig grunt [aggressive vocalization that sounds like a pig grunting or a deep throated cough], soft bark [characteristic aggressive vocalization], bark, scream and pull vegetation), moderate (chest-beat [beating of chest by hands], strut-stand/run [stand in front of or runs past another with a distinctive stiff leg stance], lunge [aggressively and quickly leaning forward towards another], direct charge, indirect charge, run at/over, push) and severe aggression (hit [aggressively strike/slap/push, etc another], attack [jumps on, bites, etc. another], drag [pulls another along the ground], fight [two individuals attack, bite, scream etc. at each other], bite, chase, kick). To quantify “aggression direction”, we assigned a score to each aggressive interaction, calculated by subtracting the standardized Elo-score of the aggressor from that of the recipient. The score was maximum (=1) for interactions where the aggressor was the lowest-ranking female and the recipient was the highest-ranking female; and minimum (=-1) for interactions where the aggressor was the highest-ranking female and the recipient was the lowest-ranking female. More generally, positive scores represented aggression up and negative scores represented aggression down the hierarchy (Figure 1).

**Figure 1.**
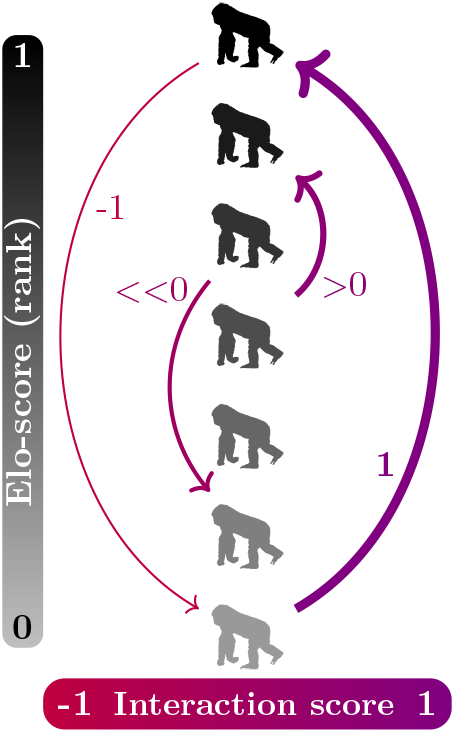
Calculation of the interaction score. The lower the rank of the aggressor and the greater the rank of the recipient, the greater the score (-1 to 1; line thickness). Arrows start from aggressors and point to recipients of aggression. Figure created using a female gorilla silhouette icon from http://phylopic.org and TikZ (TeX).

We used demographic data to estimate daily female reproductive state. On a given day, we classified as ‘pregnant’ any female that gave birth 255 days or less after that day [40], as ‘cycling’ any female that was not classified as pregnant and she had been observed mating since her last parturition and as ‘lactating’ any female with a dependent infant (based on last observation of nipple contact; [41, 42]) that was not pregnant and had not observed mating since her last parturition. Lactation is often considered more energetically demanding than pregnancy as a whole but the latest stages of pregnancy are highly energetically demanding, potentially even more than lactation [43, 44]. Thus, we differentiated between the first (1-85th day of pregnancy), second (85-170th day), and third (170-255th day) trimester of pregnancy (85 days each).

### Statistical analyses

We fitted a linear mixed effects model with a logit function to test whether females who have greater energetic needs and/or experience different social environments, direct aggression of higher or lower score (response variable, continuous, between -1 and 1; Figure 1) to other females. Given that we tested for the aggression direction and not aggression rates, the design of our analysis was independent of observation effort, and thus, we were able to use both focal and ad libitum observations. Specifically, we considered each aggressive interaction recorded during either a focal or an libitum observation as a separate data point. In our model, we fitted the following explanatory variables: aggression intensity to test if aggression of greater score is more often mild than moderate or severe; number of adult males in the group (*>*14 years old for western males; *>*12 years old for mountain males); number of females in the group; reproductive state of the aggressor (cycling, trimester of pregnancy, or lactation); and species (western or mountain). We fitted the identities of interacting females, dyad and group as random factors. Finally, we used Tukey post hoc comparisons (via the *glht* function from the *multcomp* package [45]), to perform pairwise comparisons between all reproductive states.

We ran the model in R version 4.1.2 using the function *glmmTMB* from the package *glmmTMB* [46]. We used the function *Anova* from package *car* [47] to test the significance of fixed factors and to compute 95% confidence intervals. We tested the residual distributions, using the functions *testDispersion* and *testUniformity* from package *DHARMa* version 0.4.6 [48] to validate the model. We used the base function *cor.test* and the function *check collinearity* from the *performance* package to test for correlations and multicollinearities of the explanatory variables: all VIF (variance inflation factor) values were *<*1.5 indicating no serious multicollinearities [49].

## Results

We analyzed 6871 aggressive interactions among a total of 31 adult female gorillas in the five social groups (Table 2). The average percentage of aggressive interactions directed from lower- to higher-ranking females across groups was 41.8±6.7% (±SD; ATA: 47.2%; BIT: 46.3%; KYA: 36.5%; MUK: 46.1%; ORU: 32.7%). For comparison, only 16.4±4.1% of displacement/avoidance interactions that we used to infer the highly stable hierarchies (details in [28]) were directed from lower- to higher-ranking females (±SD; ATA: 15.4%; BIT: 19.2%; KYA: 12.8%; MUK: 22.1%; ORU: 12.5%). This result confirms previous evidence that aggression may not be a reliable proxy of power in gorillas or other species [29, 30, 34].

**Table 2:**
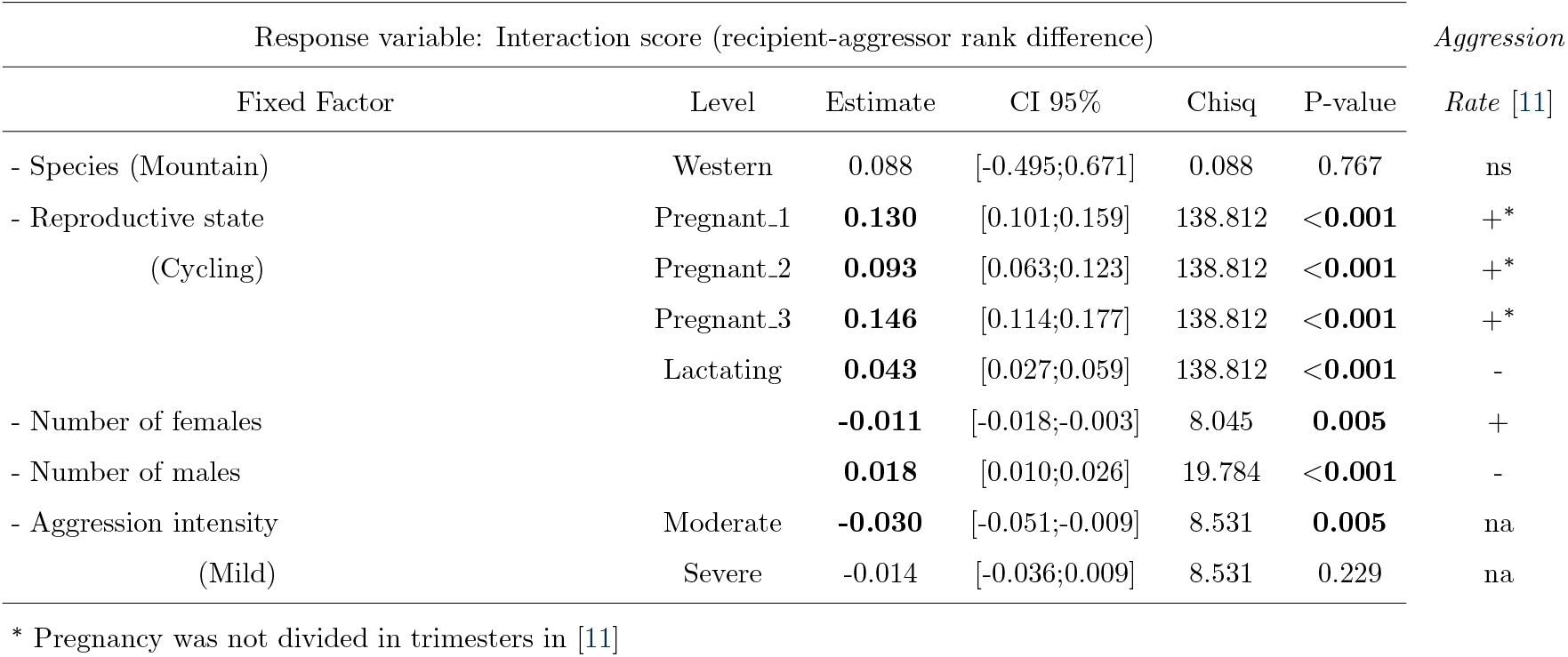
Results from the linear mixed effects model. Significant p-values appear in bold. The significance of each level of a categorical variable was evaluated against the reference level (placed in parenthesis) according to whether their confidence intervals (CI) include zero or not. ‘Pregnant n’ denotes the nth trimester of pregnancy. To highlight that aggression rates can increase due to increase in interactions of different score, we also include the effect of some of the tested variables on overall adult female aggression rates, based on results of linear mixed effects models from [11] on the right of the table. ‘ns’: non-significant correlation; ‘+’: positive correlation; ‘-’: negative correlation; ‘na’: not tested (see [11] for details).

| Response variable: Interaction score (recipient-aggressor rank difference) |  |  |  |  |  | *Aggression* |
| --- | --- | --- | --- | --- | --- | --- |
| Fixed Factor | Level | Estimate | CI 95% | Chisq | P-value | *Rate* [11] |
| - Species (Mountain) | Western | 0.088 | [-0.495;0.671] | 0.088 | 0.767 | ns |
| - Reproductive state<br>(Cycling) | Pregnant_1 | **0.130** | [0.101;0.159] | 138.812 | **<0.001** | + |
|  | Pregnant_2 | **0.093** | [0.063;0.123] | 138.812 | **<0.001** | + |
|  | Pregnant_3 | **0.146** | [0.114;0.177] | 138.812 | **<0.001** | + |
|  | Lactating | **0.043** | [0.027;0.059] | 138.812 | **<0.001** | - |
| - Number of females |  | **-0.011** | [-0.018;-0.003] | 8.045 | **0.005** | + |
| - Number of males |  | **0.018** | [0.010;0.026] | 19.784 | **<0.001** | - |
| - Aggression intensity<br>(Mild) | Moderate | **-0.030** | [-0.051;-0.009] | 8.531 | **0.005** | na |
|  | Severe | -0.014 | [-0.036;0.009] | 8.531 | 0.229 | na |

\* Pregnancy was not divided in trimesters in [11]

Aggression from lower- to higher-ranking females was 85% mild, 9% moderate and 6% severe while aggression from higher- to lower-ranking females was 82% mild, 10% moderate and 8% severe. Generally, aggression of different intensity showed similar distributions and all aggression was most common from higher- to lower-ranking females close in rank (-0.5 *>* score *>* 0; Figure 2). The interactions of mild aggression were of greater score (recipient-aggressor rank difference) than interactions of moderate aggression, meaning that females were more likely to use mild rather than moderate aggression against more powerful rivals, but the score difference between interactions of severe and moderate or mild aggression was not significant (Table 2).

**Figure 2.**
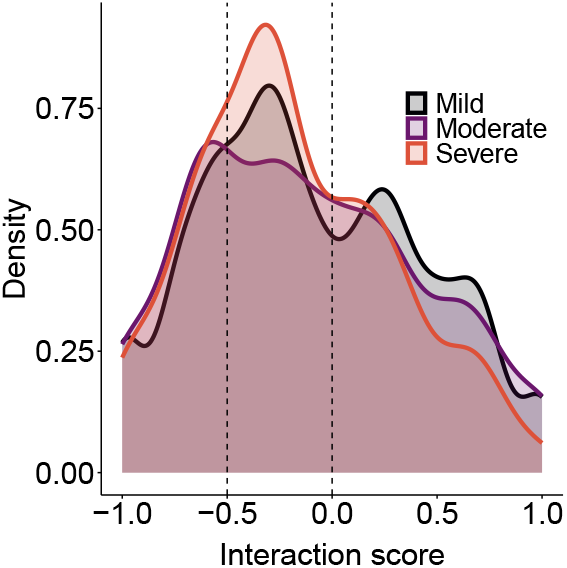
Distribution of interaction score (recipient-aggressor rank difference): density of the mild, moderate and severe aggression as a function of the interaction score. Positive scores represented aggression up and negative scores represented aggression down the hierarchy.

Female gorillas directed aggression of greater score when there were more males in the group and when there were fewer females in the group (Figure 3; Table 2). When we ran our analysis testing for group size (number of weaned individuals in the group), instead of the numbers of males and females, its influence on interaction score was not significant (estimate=-0.001, p-value=0.682). Females in the third trimester of pregnancy directed the aggression of the highest score, females in any pregnancy stage directed aggression of greater score than lactating and cycling females, and lactating females directed aggression of greater score than cycling females, likely highlighting an effect of energetic needs on aggression heuristics (Figure 3; Table 2). Post hoc pairwise comparisons showed significant differences between all reproductive states, apart from the difference of the first and the other two trimesters of pregnancy (Table 3; Figure 3). When we ran our analysis merging pregnancy trimesters into one category, we found pregnant females to direct aggression of significantly greater score than lactating and cycling females, and lactating females to direct aggression of significantly greater score than cycling females (not shown). Finally, our results did not show any significant difference between aggression score within the one western and four mountain gorillas groups.

**Table 3:** Post hoc comparisons of the different reproductive state. Significant p-values appear in bold. Cycl: cycling; P n: nth pregnancy trimester; Lact: lactating.

| Comparison | Estimate | Std.Error | Z value | p value |
| --- | --- | --- | --- | --- |
| P_1 - Cycl | **0.130** | 0.015 | 8.681 | **0.000** |
| P_2 - Cycl | **0.093** | 0.015 | 6.071 | **0.000** |
| P_3 - Cycl | **0.146** | 0.016 | 9.147 | **0.000** |
| Lact - Cycl | **0.043** | 0.008 | 5.236 | **0.000** |
| P_1 - P_2 | -0.037 | 0.019 | -1.965 | 0.267 |
| P_1 - P_3 | 0.016 | 0.019 | 0.812 | 0.922 |
| Lact - P_1 | **-0.087** | 0.015 | -5.888 | **0.000** |
| P_2 - P_3 | **0.052** | 0.019 | 2.721 | **0.047** |
| Lact - P_2 | **-0.050** | 0.015 | -3.368 | **0.006** |
| Lact - P_3 | **-0.103** | 0.015 | -6.715 | **0.000** |

**Figure 3.**
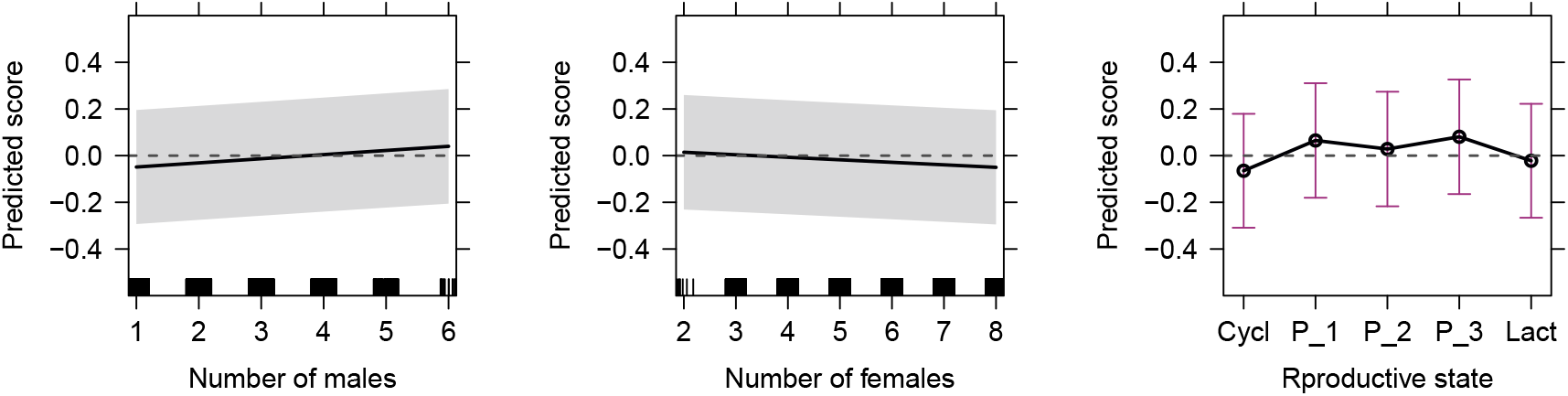
Predicted interaction score (recipient-aggressor rank difference) as a function of the explanatory variables of the linear mixed effects model, with a significant effect: number of adult males in the group, number of adult females in the group and aggressor’s reproductive state (Cycl: cycling; P n: nth pregnancy trimester; Lact: lactating). Shaded areas and whiskers show 95% confidence intervals. We created the figure using R package *effects* [50]. Positive scores represented aggression up and negative scores represented aggression down the hierarchy.

Notably, a positive (or negative) correlation of a predictor with the interaction score does not necessarily represent a shift from aggression towards females lower-ranking than the aggressor to aggression towards females higher-ranking than the aggressor (or vice versa) – but, more generally, it represents a shift of aggression towards more (or less) powerful females independently of the rank relationship to the aggressor. For example, females in the second trimester of pregnancy direct most aggression towards rivals higher-ranking than themselves; yet they direct aggression of significantly lower score than females in the third trimester of pregnancy, that is, the latter direct more aggression to females even more higher-ranking than those during second trimester (Figure 3; Table 2).

## Discussion

Most aggression was directed from higher to lower-ranking females, usually close in rank, supporting the hypothesis that individuals commonly use aggression to reinforce their status. However, approximately 42% of aggression was directed from lower- to higher-ranking females, which is even greater than previous estimations in gorillas (34% [29] & 25% [30]) and also greater than in many other animals [8]. Our results suggest that such aggression towards more powerful rivals reflects competitive incentives influenced by the social environment and driven by circumstantial needs. This interpretation is in line with previous observations showing that female gorillas are usually unable to improve their ranks through active competition, as they form highly stable hierarchical relationships [28], that is, aggression is unlikely to be used for challenginf the hierarchy. Importantly, female gorillas are more likely to respond aggressively (‘retaliate’) than submissively to aggression from other females [29], meaning that aggression involves some risk for the aggressor, especially when it targets more powerful groupmates: greater interaction score, greater risk (the greater the relative rank of the recipient, the greater the risk for the aggressor). Thus, our results further suggest that the social environment and circumstantial needs may influence individual decisions to engage in risky behaviours.

Female gorillas appeared to direct aggression of greater score when there were more males in the group, potentially supporting our interpretation that male support or protection [15, 29] provides females with an environment to take greater risks. Males may support lower-ranking females in order to decrease competitive inequities among females (e.g., to prevent low ranking female emigration from their group; [51]) and high-ranking females might hesitate to retaliate to aggressors and escalate a contest if more males are adjacent to the aggressors or are simply present in the group and can intervene. Interestingly, when females have access to fewer males, they generally exhibit greater aggression rates towards other females ([11]; Table 2, column ‘Aggression rates’), potentially competing for male protection per se; but once they have access to more males (and male protection), they appear to direct more aggression to more powerful female rivals.

In contrast to the number of adult males, the number of adult females in the group was negatively correlated with interaction score, that is, female gorillas directed aggression to lower-ranking, less powerful, females when there were more females in the group. Hence, the previously observed increase in aggression rates in groups with more females ([11]; Table 2, column ‘Aggression rates’) likely pertains predominantly to aggression from more to less powerful rivals. Our result may reflect that females preferably target less powerful rivals when they have access to a larger pool of competitors to choose from, similar to humans ([52]; see also risk-sensitivity theory in introduction). Alternatively, the increased competition due to a larger number of competitors may prompt higher-ranking females to reinforce their status more than it prompts lower-ranking females to target higher-ranking ones in order to access resources. Overall, the combination of our present and previous results [11] showing the influence of the number of males and females on both female aggression rates and aggression direction confirms that non-human animals can adapt their aggression patterns according to the social context or the available social information (see also [53]).

Aggression score was also influenced by energetic needs: female gorillas in the most energetically demanding reproductive states directed aggression to more powerful females (greater aggression score/ recipient-aggressor rank difference). Females in the last and most energetically demanding stage of pregnancy directed aggression of greater score than all other females (this difference was significant for all reproductive states except females in the first trimester of pregnancy), females in any stage of pregnancy directed aggression of greater score than lactating and cycling females, and lactating females directed aggression of greater score than cycling females. While lactating females can potentially have greater energetic needs than pregnant females, they might direct aggression of lower score than pregnant females because they show lower risk-tolerance in aggression towards higher-ranking groupmates in order to protect their dependent infants, reminiscent of risk-avoidance behaviours of lactating female African wild dogs (*Lycaon pictus*; [54]) and black bears (*Ursus americanus*; [55]). This interpretation is consistent with previous results showing that lactating females show the lowest aggression rates ([11]; see also column ‘Aggression rates’ in Table 2). Yet, lactating females directed aggression of greater score than cycling females, despite the fact that they exhibit lower aggression rates than cycling females ([11]; Table 2, column ‘Aggression rates’). Lactating females have greater proximity to alpha males [56, 57], potentially offering them greater access to resources with no need for direct female-female competition (low aggression rates), but it might also buffer female-female aggression risk if males mediate/intervene in female aggressive interactions [15, 29], allowing lactating females to direct aggression to generally more powerful rivals (high aggression score), even if this aggression is infrequent.

Our study suggests that aggression heuristics depend on the social environment and individual needs, meaning that the variation in heuristics observed within or between species [8] at least partially reflects differences in the conditions that specific animal populations or individuals experience at different time points. Our study also adds to behavioural observations from several species suggesting that individuals with greater needs might engage in more risky behaviors [58–60], including inter-individual competition [61–63]. Accordingly, it improves our understanding on the evolution of risk-taking in hominids, including (aggressive) competition in humans where individuals unsuccessful in economic or mating competition may exhibit risky aggressive behaviours [64–68]. Finally, our results may provide some insights regarding the evolution of more egalitarian or despotic societies of other species: if certain social factors or individual conditions can drive shifts in aggression up or down the hierarchy, then they have the potential to flatten or reinforce the hierarchy, respectively.

## Ethics

We followed the regulations of Agence Nationale des Parcs Nationaux and the Centre National de la Recherche Scientifique et Technique of Gabon as well as the regulations of Uganda Wildlife Authority and the Uganda National Council of Science and Technology in Uganda. Ethical clearance was given by the Max Planck Society.

## Data availability

The data and code necessary to replicate this study are available at https://gitlab.com/nksmt/GorillaHeuristics.

## Authors’ contributions

N.S.: conceptualization, data curation, formal analysis, investigation, methodology, writing—original draft and writing—review and editing; M.M.R.: data curation, funding acquisition, investigation, methodology, writing—review and editing.

## Acknowledgements

We thank all staff who assisted with data collection, project management, and logistical support in both study sites (see https://www.eva.mpg.de/primate-behavior-and-evolution/research-groups/gorilla-group/). We thank also Andrew M. Robbins and Fernando Colchero for the useful feedback in the course of this project as well as Jack L. Richardson and Christopher Young for long-term database management. We thank the Uganda Wildlife Authority and the Ugandan National Council for Science and Technology for permission to work in Bwindi Impenetrable National Park, Uganda as well as the Institute of Tropical Forest Conservation for logistical support in Bwindi. We thank the Agence Nationale des Parcs Nationaux and the Centre National de la Recherche Scientifique et Technique of Gabon for permission to work in Loango and for the help in project management.

